# PKC regulates the production of fibroblast growth factor 23 (FGF23)

**DOI:** 10.1101/520254

**Authors:** Ludmilla Bär, Philipp Hase, Michael Föller

**Affiliations:** Institute of Agricultural and Nutritional Sciences, Martin Luther University Halle-Wittenberg, D-06120 Halle (Saale), Germany; Institute of Physiology, University of Hohenheim, D-70599 Stuttgart, Germany

## Abstract

Serine/threonine protein kinase C (PKC) is activated by diacylglycerol that is released from membrane lipids by phospholipase C in response to activation of G protein-coupled receptors or receptor tyrosine kinases. PKC isoforms are particularly relevant for proliferation and differentiation of cells including osteoblasts. Osteoblasts/osteocytes produce fibroblast growth factor 23 (FGF23), a hormone regulating renal phosphate and vitamin D handling. PKC activates NFκB, a transcription factor complex controlling FGF23 expression. Here, we analyzed the impact of PKC on FGF23 synthesis. *Fgf23* expression was analyzed by qRT-PCR in UMR106 osteoblastlike cells and in IDG-SW3 osteocytes. Phorbol ester 12-O-tetradecanoylphorbol-13-acetate (PMA), a PKC activator, up-regulated *Fgf23* expression. In contrast, PKC inhibitors calphostin C, Gö6976, sotrastaurin and ruboxistaurin supressed *Fgf23* gene expression. NFκB inhibitor withaferin A abolished the stimulatory effect of PMA on *Fgf23.* PKC is a powerful regulator of FGF23 synthesis, an effect which is at least partly mediated by NFκB.

## Introduction

Protein kinase C (PKC) isoforms are related serine/threonine kinases probably expressed in all cell types. Classically, PKC activity is induced upon stimulation of various G_q_ protein-coupled receptors and growth factor receptor tyrosine kinases [1]. Three classes of PKC isoforms can be distinguished: classical PKC (cPKC) isoforms are activated by both, diacylglycerol (DAG) and an increase in the intracellular Ca^2+^ concentration whereas novel PKC (nPKC) isoforms require only DAG, and atypical PKC (aPKC) isoforms are induced by other mechanism [2]. The classical activation is dependent on phospholipase Cβ or Cγ-mediated breakdown of membrane phosphatidylinositol 4,5-bisphosphate (PIP_2_) yielding inositol 1,4,5-trisphosphate (IP_3_) and DAG. IP3 binds the IP3 receptor releasing Ca^2+^ from the endoplasmic reticulum (ER), while membrane-bound DAG activates PKC [1].

PKC is crucial for most cellular responses including the regulation of gene expression, cell migration, proliferation, differentiation, and apoptosis [3]. Moreover, PKC is implicated in the pathophysiology of frequent disorders such as heart failure, diabetes, Alzheimer and Parkinson disease, as well as inflammatory and immune disorders [3]. PKC is particularly relevant for various malignancies, owing to its tumor and metastasis-promoting properties [3]. Plant-derived phorbol esters are potent carcinogens that are effective through stimulating PKC activity [3].

Inflammation [4,5], renal and cardiovascular disease [6–8] are major triggers of the production of fibroblast growth factor 23 (FGF23), a proteohormone mainly produced in the bone [9] and implicated in the regulation of phosphate reabsorption and 1,25(OH)_2_D_3_ (active vitamin D [10]) formation in the kidney [11]. The FGF23-mediated inhibition of renal phosphate transporter NapiIIa and CYP27B1, the key enzyme for the generation of 1,25(OH)_2_D_3_, is dependent on Klotho, a transmembrane protein [9]. The induction of left ventricular hypertrophy is, however, solely mediated by FGF23 without the involvement of Klotho [8,12,13], whereas vitamin D partly overcomes the FGF23 effect on cardiac hypertrophy [14]. FGF23 also impacts on neutrophil recruitment [15], erythropoiesis [16,17], or hepatic cytokine secretion [18] in a paracrine manner.

Klotho or FGF23 deficiency results in rapid aging, a very short life span, and multiple age-associated diseases affecting most organs and tissues [11,19]. This dramatic phenotype of Klotho or FGF23 null mice is almost completely rescued by a phosphate-or vitamin D-deficient diet [20,21], pointing to the predominant role of calcification in the pathophysiology of Klotho or FGF23 deficiency [22]. Apart from its endocrine and paracrine effects that are still incompletely understood, FGF23 is a putative disease biomarker [6,23]. The plasma level of FGF23 correlates well with progression of chronic kidney disease (CKD) and is a very sensitive marker that is elevated even before onset of hyperphosphatemia or hyperparathyroidism [24,25]. Further, FGF23 levels are elevated in acute kidney injury [26,27]. Whether and to which extent FGF23 not only indicates disease but actively contributes to disease progression, as shown for the heart, remains unclear.

The identification of molecular regulators of FGF23 production is of high interest and relevance. Known regulators include PTH [28], 1,25(OH)_2_D_3_ [29], phosphate [30,31], inflammatory cytokines and factors such as TNFα [32], IL-1/6 [4,33–35], or NFκB [33,36], TGFβ [37], AMP-dependent protein kinase (AMPK) [38] or insulin-dependent PI3 kinase signaling [39].

The present study explored the contribution of PKC signaling to the production of FGF23 in bone cells.

## Materials and Methods

### Cell culture

Cell culture and experiments with UMR106 rat osteoblast-like cells were conducted as described before [39]. Briefly, cells were cultured in DMEM high-glucose medium containing 10% FBS and penicillin-streptomycin at 37°C and 5% CO_2_.

IDG-SW3 bone cells were cultured as described earlier [40]. Briefly, nondifferentiated cells were kept at 33°C in AlphaMEM medium (with L-glutamine and deoxyribonucleosides) containing 10% FBS, penicillin-streptomycin and interferon-gamma (INF-γ; 50 U/ml). For differentiation, cells were plated on collagen-coated dishes at 37°C in medium with 50 μg/ml ascorbic acid and 4 mM β-glycerophosphate but without INF-γ. All reagents were from ThermoFisher unless indicated.

IDG-SW3 osteocytes were used after 35 days of differentiation, and UMR106 cells were pretreated with 100 nM 1,25(OH)_2_D_3_ (Tocris, Wiesbaden-Nordenstadt, Germany) for 24h before the experiment. Cells were then incubated with activator Phorbol ester 12-O-tetradecanoylphorbol-13-acetate (PMA; Sigma, Schnelldorf, Germany; 0.1 μM; 6 h) with or without 1 μM PKC inhibitors Calphostin C (Tocris), Gö6976 (Tocris), sotrastaurin (Selleckchem, München, Germany), ruboxistaurin (Selleckchem), or NFκB inhibitor withaferin A (Tocris; 0.5 μM), or with vehicle only for another 24 h.

### Expression analysis

Total RNA was extracted using peqGOLD TriFast reagent (VWR, Dresden, Germany). Complementary DNA (cDNA) was synthesized from 1.2 μg RNA using random primers and the GoScript™ Reverse Transcription System (Promega, Mannheim, Germany) at 25°C for 5 min, 42°C for 1 h, and 70°C for 15 min.

The expression profile of PKC isoforms in UMR106 cells was studied by RT-PCR using the GoTaq Green Master Mix (Promega) and the primers listed in Table 1. For the PCR, 2 μl synthesized cDNA were used. Settings were: 95°C for 3 min, followed by 35 cycles of 95°C for 30 s, 60°C for 30s, 72°C for 45s. PCR products were loaded on a 1.5% agarose-gel and visualized by Midori Green (Biozym, Hessisch Oldendorf, Germany), and a 100 bp DNA ladder (Jena Bioscience, Jena, Germany) was used as a size marker.

**Fig 1:**
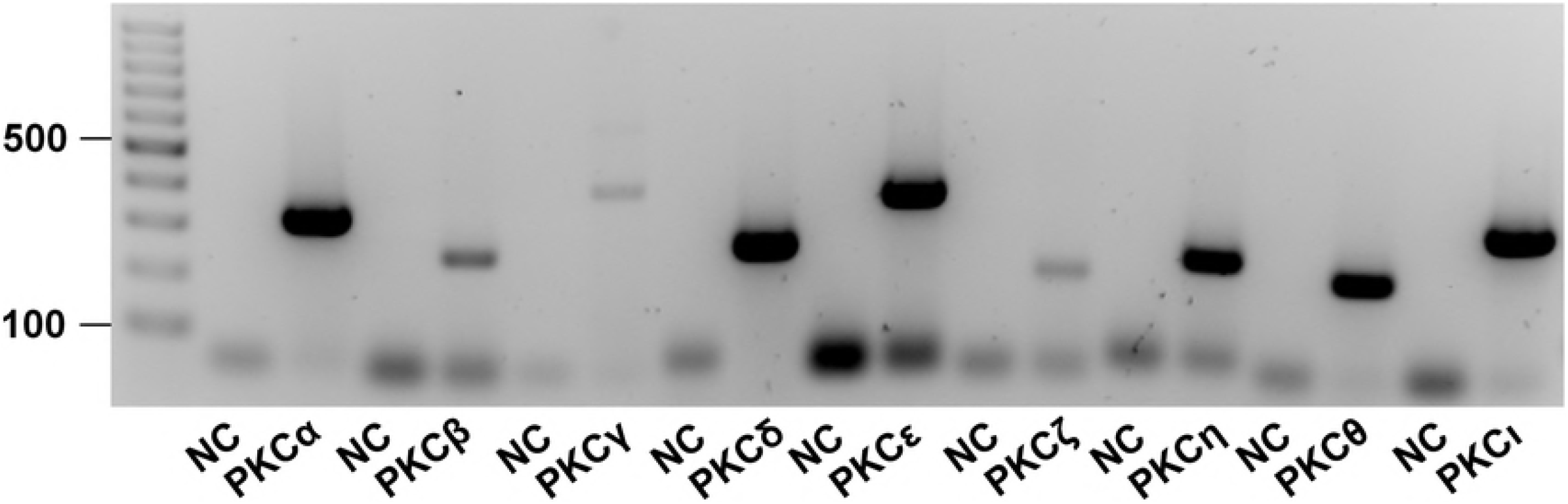
Expression of PKC isoforms in UMR106 osteoblast-like cells. Original agarose gel photo showing PKCα, -β, -γ, -δ, -ε, -ζ, -η, -θ or -ι specific cDNA in UMR106 cells. NC: non-template control

**Table 1:**
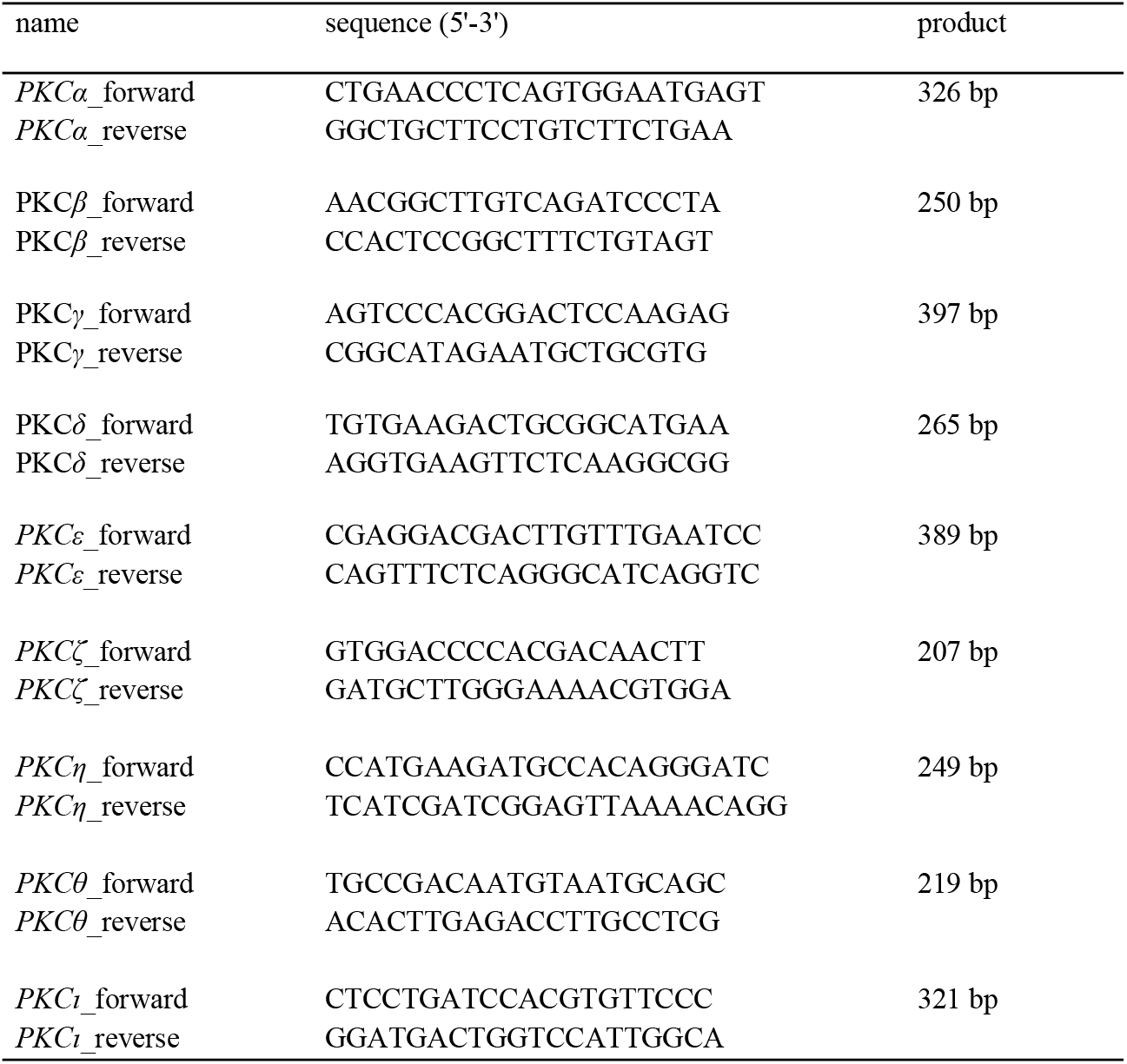
Primer sequences.

### qRT-PCR

Relative expression levels of *Fgf23* were determined by qRT-PCR using 2 μl synthesized cDNA and the GoTaq qPCR Master Mix (Promega) on a Rotor-Gene Q (Qiagen, Hilden, Germany). PCR conditions were 95°C for 3 min, followed by 35 cycles of 95°C for 10 s, 58°C for 30 s and 72°C for 45 s. After normalization to *Tbp* (TATA box-binding protein) expression, relative quantification of gene expression based on the double-delta Ct (threshold cycle) analysis was carried out.

Primers used:

*Tbp*

F: ACTCCTGCCACACCAGCC

R: GGTCAAGTTTACAGCCAAGATTCA

*Fgf23*

F: TGGCCATGTAGACGGAACAC R: GGCCCCTATTATCACTACGGAG

### Statistics

Arithmetic means ± SEM were calculated, and *n* represents the number of independent experiments. Comparisons of two groups were made by unpaired Student’s t test, and for more than two groups, comparisons were calculated via one-way ANOVA, followed by Tukey’s or Dunnett’s multiple comparison tests, using GraphPad Prism. Differences were considered significant if p < 0.05.

## Results

The relevance of PKC activity for the synthesis of FGF23 was studied in UMR106 osteoblast-like cells and IDG-SW3 osteocytes. First, the expression of PKC isoforms was explored by RT-PCR. As demonstrated in Fig 1, mRNA specific for PKCα, PKCδ, PKCε, PKCη, PKCθ, and PKCι could readily be detected. The bands indicating the abundance of PKCβ, PKCγ, PKCζ mRNA in UMR106 cells were weaker albeit detectable.

Phorbol ester 12-O-tetradecanoylphorbol-13-acetate (PMA) is a potent activator of PKC [3]. We treated UMR106 cells with and without PMA and determined *Fgf23* transcripts by qRT-PCR. PMA treatment significantly up-regulated the abundance of *Fgf23* mRNA (Fig 2A). Similar to osteoblasts, PKC activation with PMA enhanced *Fgf23* gene expression in IDG-SW3 osteocytes (Fig 2B). These results suggest that PKC activity drives *Fgf23* gene expression in osteoblasts and osteocytes.

**Fig 2:**
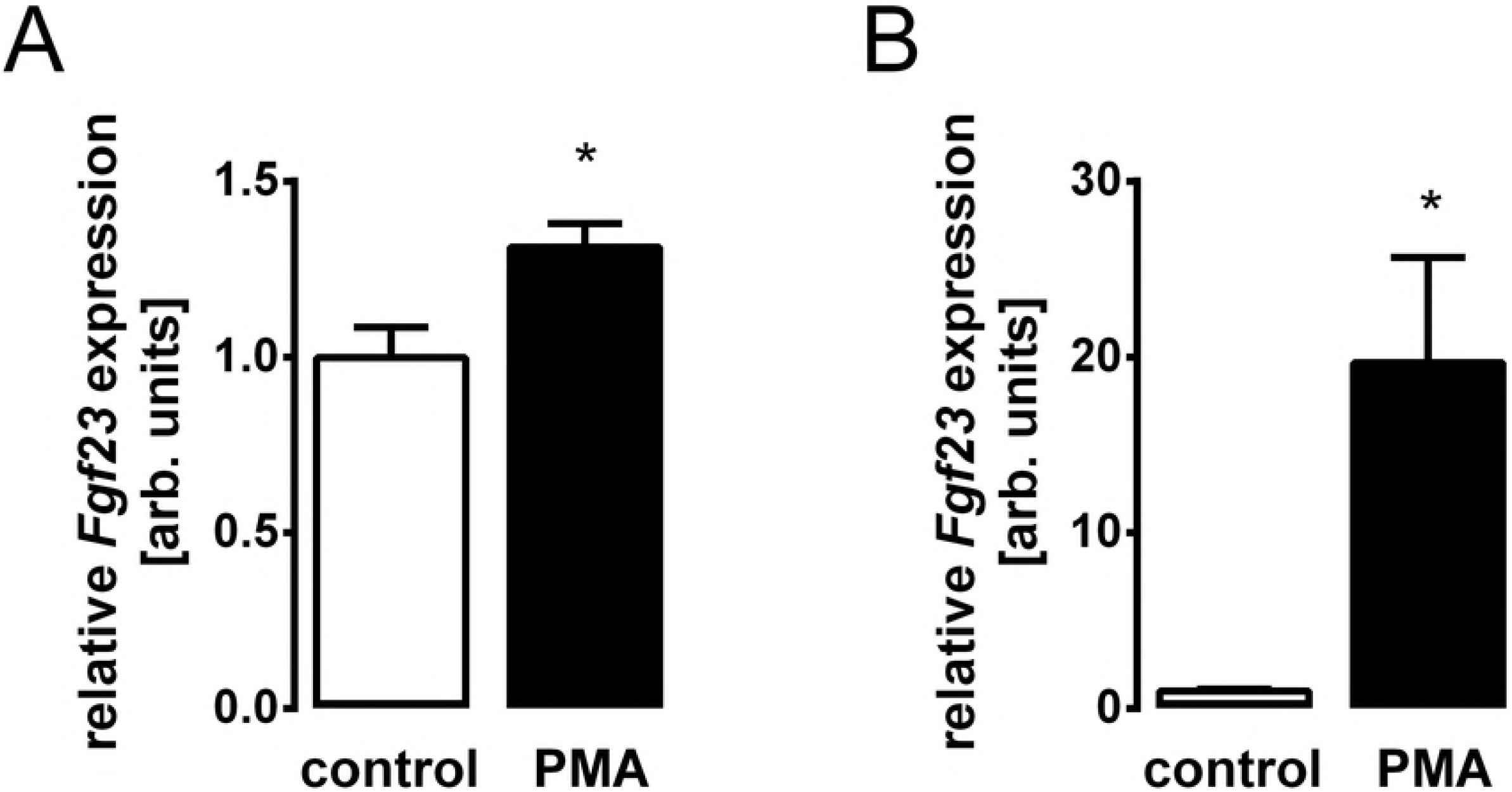
PKC activator PMA induces *Fgf23* gene expression in UMR106 osteoblast-like cells and in IDG-SW3 osteocytes. Arithmetic means ± SEM (n=6) of relative *Fgf23* mRNA abundance normalized to *Tbp* in UMR106 osteoblast-like cells **(A)** or IDG-SW3 osteocytes **(B)** incubated without (white bars) or with (black bars) 0.1 μM PKC activator PMA. * *p* < 0.05 indicates significant difference. arb., arbitrary

Our next series of experiments tested whether inhibition of PKC interferes with *Fgf23* gene expression. To this end, UMR106 cells were exposed to PKC inhibitors. As demonstrated in Fig 3, PKC inhibitor calphostin C (Fig 3A) and also PKCα/β inhibitor Gö6976 (Fig 3B) significantly and dose-dependently down-regulated *Fgf23* gene expression in UMR106 cells. Thus, PKC is a stimulator of *Fgf23* gene expression.

**Fig 3:**
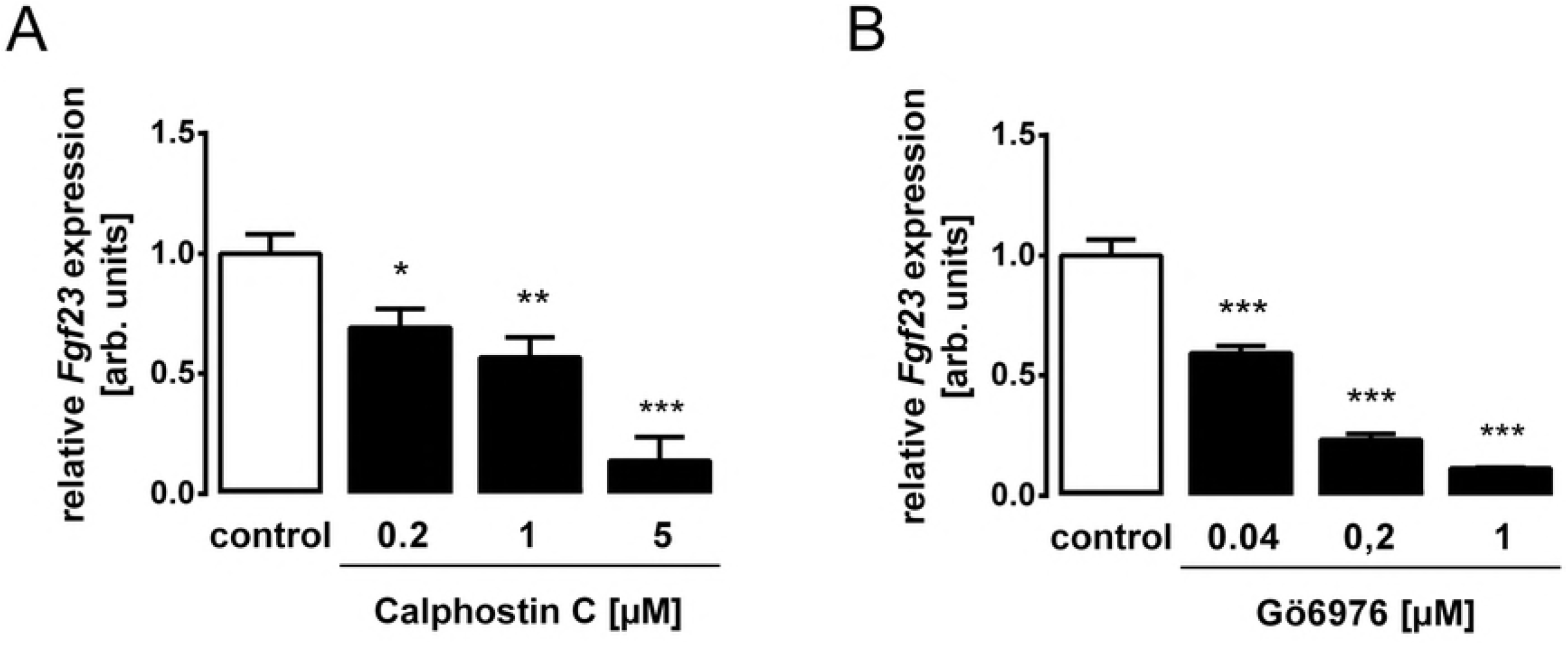
PKC inhibitors Calphostin C and Gö6976 decrease *Fgf23* transcript levels in UMR106 osteoblast cells. The bar diagrams represent the relative mRNA abundance of *Fgf23* in UMR106 cells incubated without or with PKC inhibitors Calphostin C **(A)** or Gö6976 **(B)** at the indicated concentrations. Gene expression was normalized to *Tbp* as a housekeeping gene, and the values are expressed as arithmetic means ± SEM (n = 6). \**p* < 0.05, \*\**p* < 0.01, and \*\*\**p* < 0.001 indicate significant difference.

Furthermore, we investigated whether PMA-stimulated *Fgf23* gene expression is indeed dependent on PKC activity using UMR106 and IDG-SW3 cells. As demonstrated in Fig 4, the PMA effect on *Fgf23* gene expression was completely abrogated by PKC inhibitor Gö6976 in UMR106 osteoblast-like cells (Fig 4A) and in IDG-SWR3 osteocytes (Fig 4B), and also by PKC inhibitors sotrastaurin (Fig 4C) and ruboxistaurin (Fig 4D) in UMR106 cells.

**Fig 4:**
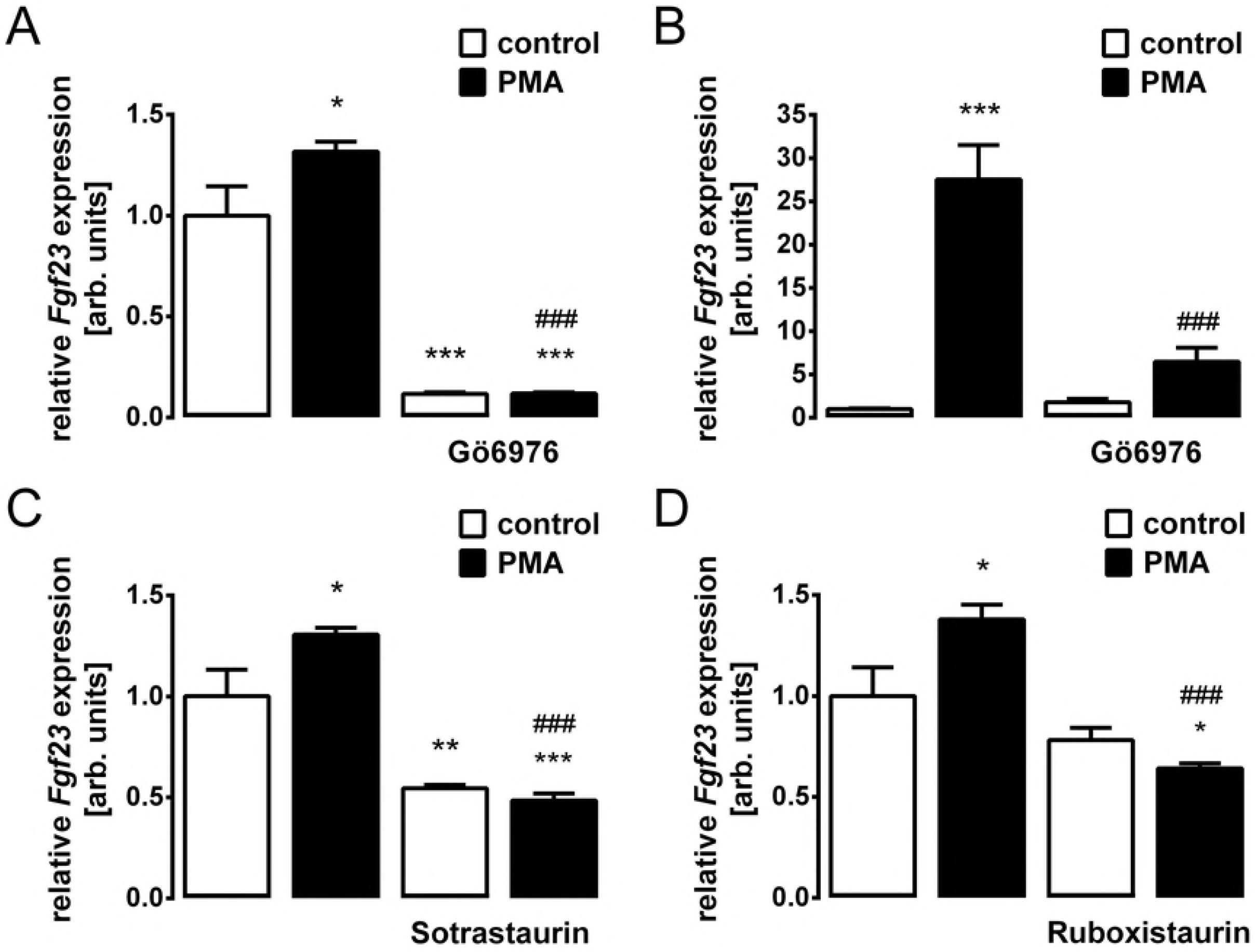
PKC inhibition abrogates the PMA-induced increase in *Fgf23* gene expression in UMR106 osteoblast-like cells and in IDG-SW3 osteocytes. Relative *Fgf23* transcript levels in UMR106 cells **(A,C,D)** or in IDG-SW3 cells **(B)** incubated without or with PMA (0.1 μM, **A-D**) in the absence and presence of PKCα/β inhibitor Gö6976 (1 μM, **A,B**), pan PKC inhibitor Sotrastaurin (1 μM, **C**) or PKCβ inhibitor Ruboxistaurin (1 μM, **D**). Gene expression was normalized to *Tbp* as a housekeeping gene, and the values are expressed as arithmetic means ± SEM (n = 6). *p < 0.05, \*\**p* < 0.01, and \*\*\**p* < 0.001 indicate significant difference from vehicle (first bar). *###p* < 0.001 indicates significant difference from the absence of PKC inhibitor (second bar vs. fourth bar). arb., arbitrary.

Finally, we sought to identify the mechanism of PKC-dependent FGF23 regulation. Since PKC is an activator of NFκB [41], a known regulator of FGF23, we treated UMR106 cells with and without NFκB inhibitor withaferin A in the absence and presence of PMA and determined *Fgf23* gene expression. As demonstrated in Fig 5, the PMA effect was indeed abrogated by withaferin A. Hence, PKC was, at least in part, effective through NFκB activity.

**Fig 5:**
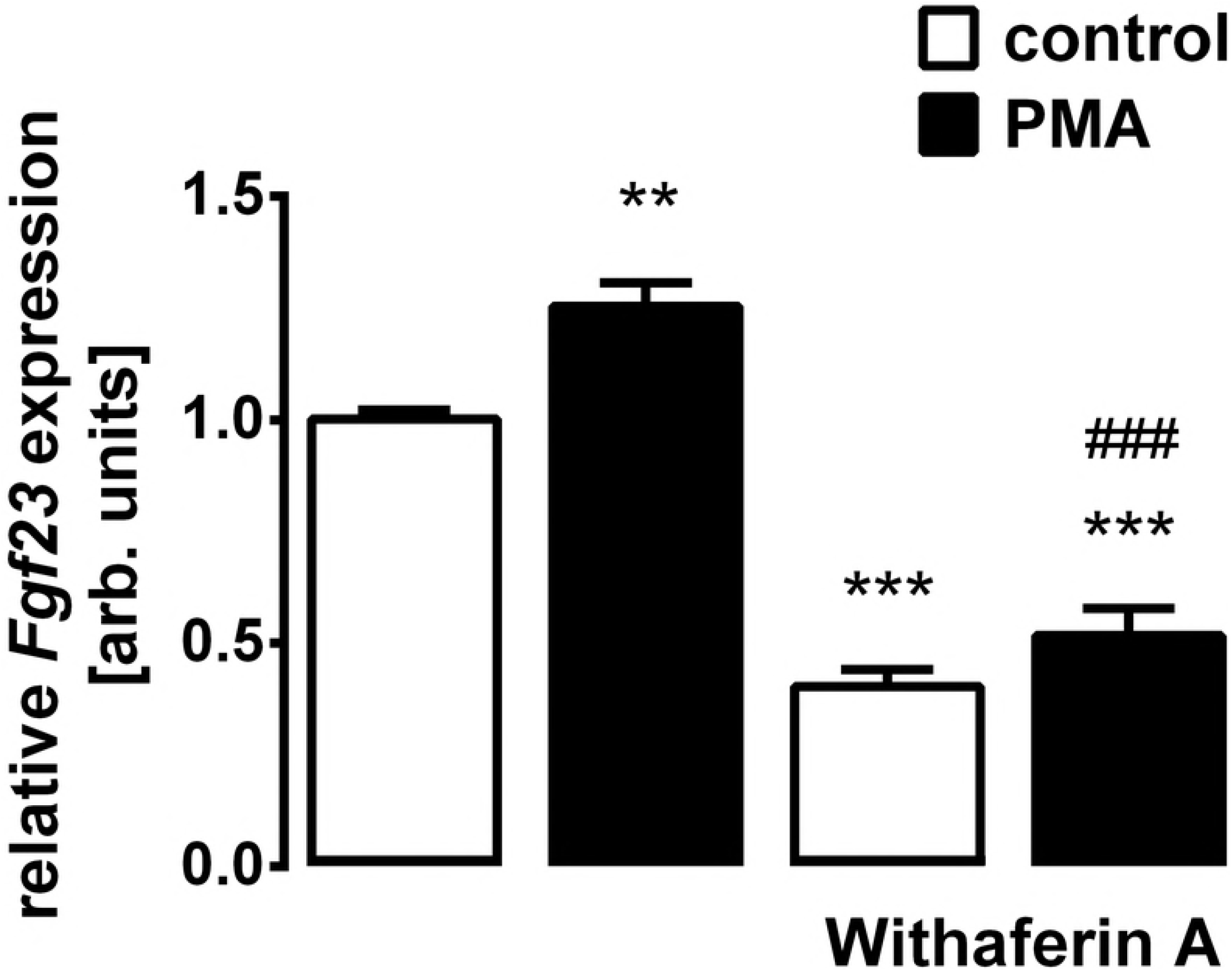
The PKC effect on *Fgf23* gene expression is dependent on NFκB. Relative *Fgf23* transcript levels in UMR106 cells incubated without or with PMA (0.1 μM) in the absence and presence of NFκB inhibitor Withaferin A (0.5 μM). Gene expression was normalized to *Tbp* as a housekeeping gene, and the values are expressed as arithmetic means ± SEM (n = 6). \*\**p* < 0.01, and \*\*\**p* < 0.001 indicate significant difference from vehicle (first bar). ###*p* < 0.001 indicates significant difference from the absence of withaferin A (second bar vs. fourth bar). arb., arbitrary.

## Discussion

Our study discloses PKC as a novel regulator of FGF23 production. We provide experimental evidence that activation of PKC induces and inhibition of PKC suppresses *Fgf23* gene expression: Four different PKC inhibitors were similarly capable of completely blocking PMA-induced *Fgf23* expression. This result unequivocally demonstrates the powerful role of PKC in regulating FGF23 production.

FGF23 is mainly produced by osteoblasts/osteocytes in the bone [9]. The differentiation of osteoblasts is driven by transcription factor MSX2 [42]. PKC enhances the proliferation of osteoblasts [43]. In contrast, PKC inhibits the differentiation of osteoblasts by targeting MSX2 [44] whereas FGF23 induces MSX2. [45]. Transcriptional activity of Runx2, also implicated in osteoblast differentiation, is enhanced by both PKC [46] and FGF23 [47].

We could show that four different pharmacological PKC inhibitors potently suppressed *Fgf23* gene expression. PKC inhibition has been suggested as a therapeutic approach in multiple diseases including cancer, sequelae of diabetes, cardiovascular diseases, or inflammatory disorders such as psoriasis [48]. Particularly renal and cardiovascular diseases are associated with elevated FGF23 plasma levels [6], and FGF23 not only indicates disease, but actively contributes at least to left heart hypertrophy [12,13]. Therefore, the therapeutic benefit of PKC inhibitors may at least in part also be due to their FGF23-lowering capacity.

Two different cell lines representing both osteoblasts and mature osteocytes were used in our study to decipher the effect of PKC on FGF23: PKC activation with PMA enhanced and PKC inhibition with different PKC inhibitors suppressed *Fgf23* gene expression in both UMR106 osteoblast-like cells and IDG-SW3 osteocytes. This result suggests PKC-dependent regulation of FGF23 formation as a universal mechanism for the control of this hormone.

In an attempt to identify the underlying mechanism we found that PKC-mediated up-regulation of *Fgf23* expression is sensitive to NFκB inhibition. NFκB, as a p65/p50 dimer, is inactivated by binding inhibitory κB (IκB). The inhibitory κB kinase (IKK) phosphorylate IκB, leading to its ubiquitination and degradation, paralleled by the release and activation of NFκB [49]. Therefore, the activation of the IKK complex plays a crucial role in the induction of NFκB, and PKC activates NFκB through this kinase [41]. Clearly, inflammation is a major trigger of FGF23 production [4,36]. In detail, NFκB up-regulates Ca^2+^ release-activated Ca^2+^ (CRAC) channel Orai1/STIM1 accomplishing store-operated Ca^2+^ entry (SOCE) and inducing the transcription of the *Fgf23* gene [36]. According to our experiment with NFκB inhibitor withaferin A, the PKC effect on FGF23 was, at least partly, dependent on NFκB pointing to the decisive role of this pro-inflammatory transcription factor complex in the regulation of FGF23.

In conclusion, our study identified PKC as a novel regulator of FGF23 production in both osteoblast and osteocytes, being at least partially effective via the NFκB signaling pathway. Therefore, the therapeutic modulation of PKC activity in chronic disease may impact on the plasma FGF23 concentration.

## Acknowledgements

The authors acknowledge the technical assistance of S. Ross and F. Reipsch.

## Author contributions

**Conceptualization:** Michael Föller.

**Formal analysis:** Ludmilla Bär.

**Funding acquisition:** Michael Föller.

**Investigation:** Ludmilla Bär, Philipp Hase.

**Supervision:** Michael Föller.

**Writing:** Ludmilla Bär, Michael Föller.

